# Biological Network Organization, Not Generic Graph Topology, Drives Graph-Based Gene Essentiality Prediction

**DOI:** 10.64898/2026.06.30.735480

**Authors:** Setareh Rahimi, Stephen Bonner, Avid Afzal, Marta Milo, Edward Morrissey, Evangelia Petsalaki

**Affiliations:** European Molecular Biology Laboratory, European Bioinformatics Institute (EMBL-EBI), Wellcome Genome Campus, Hinxton, Cambridge, UK; AI for Science Innovation, Astrazeneca, CB2 0AA, cambridge; Data Sciences and Quantitative Biology, Discovery Sciences, R&D, AstraZeneca, CB2 0AA, Cambridge, UK

**Keywords:** Gene essentiality, Graph neural networks, CRISPR screening, Protein–protein interaction networks, Network topology, Functional genomics

## Abstract

Predicting gene essentiality across cellular contexts is a central challenge in computational biology, with implications for identifying cancer vulnerabilities. Graph neural networks (GNNs) integrate molecular interaction networks with gene-level features, but it remains unclear whether their performance gains arise from biologically meaningful connectivity or generic graph structure. Here, we systematically evaluate the role of network information in gene essentiality prediction using 2,741 genes across three tissues. We compare GNNs to feature-only baselines, including multilayer perceptron (MLP) and random forest (RF) methods, under a strict gene-level 5-fold cross-validation scheme to prevent information leakage. To isolate the role of network information, we assess models on the STRING protein–protein interaction network, a degree-preserving shuffled network, and a fully randomized network, with and without network-derived features. GNNs outperform feature-only models, reducing mean squared error and improving Matthews correlation coefficient across all tissues. However, these gains depend critically on biologically structured connectivity: performance degrades substantially under randomized topology and is not preserved by degree-constrained rewiring. Network features are largely redundant when using biologically meaningful graphs, as their information is recovered through message passing, but become important when topology is uninformative. Per-gene analyses reveal uniformly low correlations across models, highlighting intrinsic limits imposed by data variability. Graph Transformer models incorporating global attention do not outperform standard GNNs, indicating that predictive signals are predominantly local. Together, these results show that predictive gains arise from biologically structured connectivity rather than generic graph topology.

## Introduction

Identifying gene dependencies, genes whose loss impairs cellular viability, is a central problem in cancer biology and precision oncology, because such dependencies can reveal therapeutic vulnerabilities (Yu et al., 2025). Large-scale RNA interference and more recently CRISPR–Cas9 loss-of-function screens have generated extensive catalogs of gene essentiality across hundreds of cancer cell lines, most notably through the Cancer Dependency Map (DepMap) initiative (Wang *et al*. 2014, Meyers *et al*. 2017, Tsherniak *et al*. 2017, Behan *et al*. 2019, Dempster *et al*. 2019). However, experimental screening remains resource-intensive and cannot be performed across all tissues and genetic contexts, motivating the development of computational approaches for in silico prediction.

A key challenge in gene essentiality prediction is its context dependence: although many genes show relatively stable dependency across cell lines, others become essential only in specific cellular or genetic contexts. As a result, relationships between molecular features and dependency phenotypes are nonlinear and multifactorial, limiting generalization to unseen genes and contexts (Gönen *et al*. 2017, Tsherniak *et al*. 2017). Furthermore, many genes exhibit only limited variation in essentiality across available cell lines, reducing the predictive signal available for modeling context-specific effects.

Computational approaches have integrated gene expression, functional annotations, and protein– protein interaction (PPI) networks, for example, through supervised regression or classification tasks, to predict gene essentiality (Seringhaus *et al*. 2006, Hart *et al*. 2014). Early work leveraged network topology, motivated by observations such as the centrality–lethality relationship (Jeong *et al*. 2001, He and Zhang 2006, Li M *et al*. 2011, Wang *et al*. 2015, Li X *et al*. 2020), but topology-based features alone are insufficient due to noise and bias in biological networks (Mahdavi and Lin 2007, Zhang, Acencio, and Lemke 2016). More recently, machine learning and deep learning models have been applied to capture nonlinear relationships in high-dimensional molecular data, although reported performance gains depend strongly on feature selection, model design, and evaluation strategy (Chen and Xu 2005, Seringhaus *et al*. 2006). In addition, increased model complexity introduces risks of overfitting, highlighting the need for controlled evaluation.

Graph-based models provide a principled framework for incorporating biological structure into these predictions through neighborhood-aware representation learning (Wu *et al*. 2020). Graph neural networks (GNNs) operate directly on interaction networks through iterative message passing between neighboring nodes (Kipf and Welling, 2017; Gilmer *et al*. 2017) and have been shown to outperform feature-only models in several settings (Schapke, Tavares, and Recamonde-Mendoza 2021), including graph-based approaches such as EPGAT (Schapke et al., 2021) and recent context-aware dependency prediction models integrating transcriptomic and interaction-network information (Yu et al., 2025; Pacini et al., 2024). However, these models also introduce increased complexity and sensitivity to architectural choices, making it difficult to attribute performance gains to specific sources of signal.

Critically, it remains unclear whether improvements from graph-based models reflect biologically meaningful interaction structure or generic properties of graph topology. Protein interaction networks exhibit characteristic topological features, including highly skewed degree distributions (Barabási and Albert, 1999), and highly connected hub proteins are more likely to be essential (Jeong et al., 2001), raising the possibility that models exploit topological shortcuts rather than biological signals. Most existing studies compare different networks or model architectures without topology-matched controls, making it difficult to disentangle biological organization from structural confounding. As a result, it remains difficult to interpret model performance and assess the extent to which predictions reflect meaningful biological signals.

Here, we present a controlled evaluation framework for assessing the contribution of graph structure to gene essentiality prediction. Using DepMap CRISPR data across three tissues; breast, lung, and large intestine, we compare GNNs with feature-only baselines under a strict gene-level cross-validation scheme that prevents information leakage. To isolate the role of network structure, we evaluate models across the STRING interaction network, a degree-preserving shuffled network, and a fully random network, and assess the contribution of network-derived node features. We further test whether incorporating both local and long-range dependencies via a graph transformer improves predictive performance beyond standard message-passing architectures.

We show that GNNs outperform feature-only models, but that these gains depend critically on biologically structured connectivity and are not preserved under topology-matched randomization. Furthermore, the inclusion of global attention mechanisms does not yield additional improvements, suggesting that the relevant predictive signal is largely local.

Through this framework, we disentangle the contributions of biological connectivity, graph topology, and feature-derived information, and assess when graph structure meaningfully improves gene essentiality prediction. To our knowledge, this is the first study to systematically separate biological signals from generic graph topology in this setting using controlled network perturbations. All code is provided open source to enable full reproducibility of all experiments.

## Methods

### Data Sources and Preprocessing

We used publicly available data from the DepMap Public 24Q2 release (Broad Institute, 2024), downloaded in August 2024. Gene essentiality targets were obtained from post-Chronos gene effect estimates (“CRISPRGeneEffect.csv”), which integrate CRISPR–Cas9 screening results across cell lines and correct for copy number effects and screen quality (Dempster *et al*. 2021). These continuous scores represent single-gene knockout effects on cellular viability, where more negative values indicate stronger essentiality. Genes with missing essentiality scores were imputed with the mean score across all cell lines.

Each training sample corresponds to a (gene, cell line) pair. Analyses were conducted separately for three tissues: breast (n = 35 cell lines), lung (n = 43), and large intestine (n = 106). No cell-line-level filtering was applied.

For each (gene, cell line) pair, features combined cell-line–specific and gene-level information. Cell-line–specific features included: (i) log2(TPM + 1) gene expression values, batch-corrected for strandedness using ComBat (Johnson et al., 2007); (ii) binary somatic mutation status derived from Mutect2 calls, where a gene was assigned 1 if it harbored at least one protein-altering variant (missense or nonsense SNV, or indel) in that cell line, and 0 otherwise including cases with no available mutation data; and (iii) shortest-path distances from each gene to cancer driver genes that are mutated in that specific cell line, computed on the STRING interaction graph using the driver gene list from the Cell Model Passports (Meer van der *et al*. 2019), with missing drivers encoded as zero.

Gene-level features (cell-line invariant) were selected to capture complementary aspects of gene function, cellular context, and network organization that have previously been associated with gene essentiality. These included shortest-path distances to all cancer driver genes on the interaction graph, reflecting proximity to known cancer-relevant pathways; binary functional annotations (kinase, transcription factor, receptor), capturing broad molecular roles; one-hot encoded subcellular location, providing information on cellular compartment and interaction context; and network topology measures (degree, closeness centrality, betweenness centrality, and PageRank), describing the position and importance of genes within the interaction network. All features were concatenated into a single node feature vector per gene. The interaction topology was treated as cell-line invariant, while cellular context was represented through cell-line–specific node features.

The gene–gene interaction graph was derived from STRING v12.0 (Szklarczyk et al., 2023), retaining interactions with combined confidence score > 800. A stringent threshold was used to prioritize highly reliable interactions while minimizing noise from low-confidence associations. The graph was treated as undirected and unweighted, with self-loops included to allow each node to retain its own feature information during message passing. From an initial set of 13,367 genes overlapping STRING and DepMap datasets, we retained all genes with essentiality standard deviation ≥ 0.2 across cell lines, which accounted for approximately 80% of the final dataset. To preserve representation of lower-variance genes, the remaining ~20% were randomly sampled from genes with standard deviations between 0.10–0.15 (5%) and 0.15–0.20 (15%). This yielded a graph of 2,741 nodes and 43,598 edges (density: 1.16%, mean degree 31.8). Variance distributions were preserved across data splits, and all models used this same graph. Analyses were restricted to 2,741 genes present in both STRING and DepMap datasets across three tissues.

Continuous features were z-score normalized for neural network models; binary features were left unchanged. Random Forest models used raw feature values. Genes with missing expression values and features with negligible variance (std ≤ 1e−5) were removed prior to analysis.

### Prediction Task and Evaluation

All models were trained to predict continuous Chronos gene effect scores. A derived binary classification task defined genes as essential if score ≤ −0.5; no separate classifier was trained, and classification metrics were computed by thresholding regression outputs post hoc.

The primary training objective was mean squared error (MSE). Hyperparameter optimization jointly minimized MSE and Matthews Correlation Coefficient (MCC) in a multi-objective framework (Optuna; Akiba *et al*. 2019). Additional metrics, Pearson and Spearman correlation, macro- and weighted-F1, balanced accuracy, precision, and recall, were computed for evaluation but not directly optimized. Metrics were computed both globally across all test genes and at the gene level. For per-gene evaluation, correlations were computed for each test gene across cell lines and subsequently averaged across all test genes.

### Gene-Level Split and Semi-Supervised Setup

We adopt a gene-level split to evaluate generalization to entirely unseen genes, which represents a more stringent setting than the conventional cell-line split (**Supplementary Text S1; Figure S1)**. Under cell-line splits, simple baselines such as gene-wise mean predictors can achieve strong performance due to limited variability in gene essentiality across cell lines. Five-fold cross-validation was performed with genes as the unit of splitting: in each fold, 20% of genes formed the held-out test set, and the remaining 80% were further divided into training (80%) and validation (20%) subsets. Splits were stratified by gene-level essentiality standard deviation to ensure proportional representation of low, intermediate, and high-variance genes across train, validation and test subsets, and were constructed independently per tissue.

In the semi-supervised setting, only training genes contributed to loss computation. Validation genes were used exclusively for model selection and early stopping, and test genes were held out for final evaluation. Performance is reported as mean ± standard deviation across five cross-validation folds.

For graph-based models, we adopt a transductive semi-supervised node-level regression setup, where message passing is performed over the full interaction graph, including both labeled (training) and unlabeled (validation and test) genes. However, loss computation and backpropagation are restricted strictly to training nodes, and no label information from validation or test genes is used during training. All nodes retain their input features throughout. Feature-only models (MLP and Random Forest) are trained exclusively on labeled genes and do not operate directly on the interaction graph. While their input vectors include graph-derived features, they do not use the graph adjacency structure for neighborhood aggregation or information propagation. This transductive formulation reflects the biological setting in which the interaction network is known a priori, while essentiality labels are available only for a subset of genes.

### Model architectures and training

All models were trained and evaluated under the identical gene-level split, loss functions, and evaluation metrics described above (**Figure 1**).

**Figure 1.**
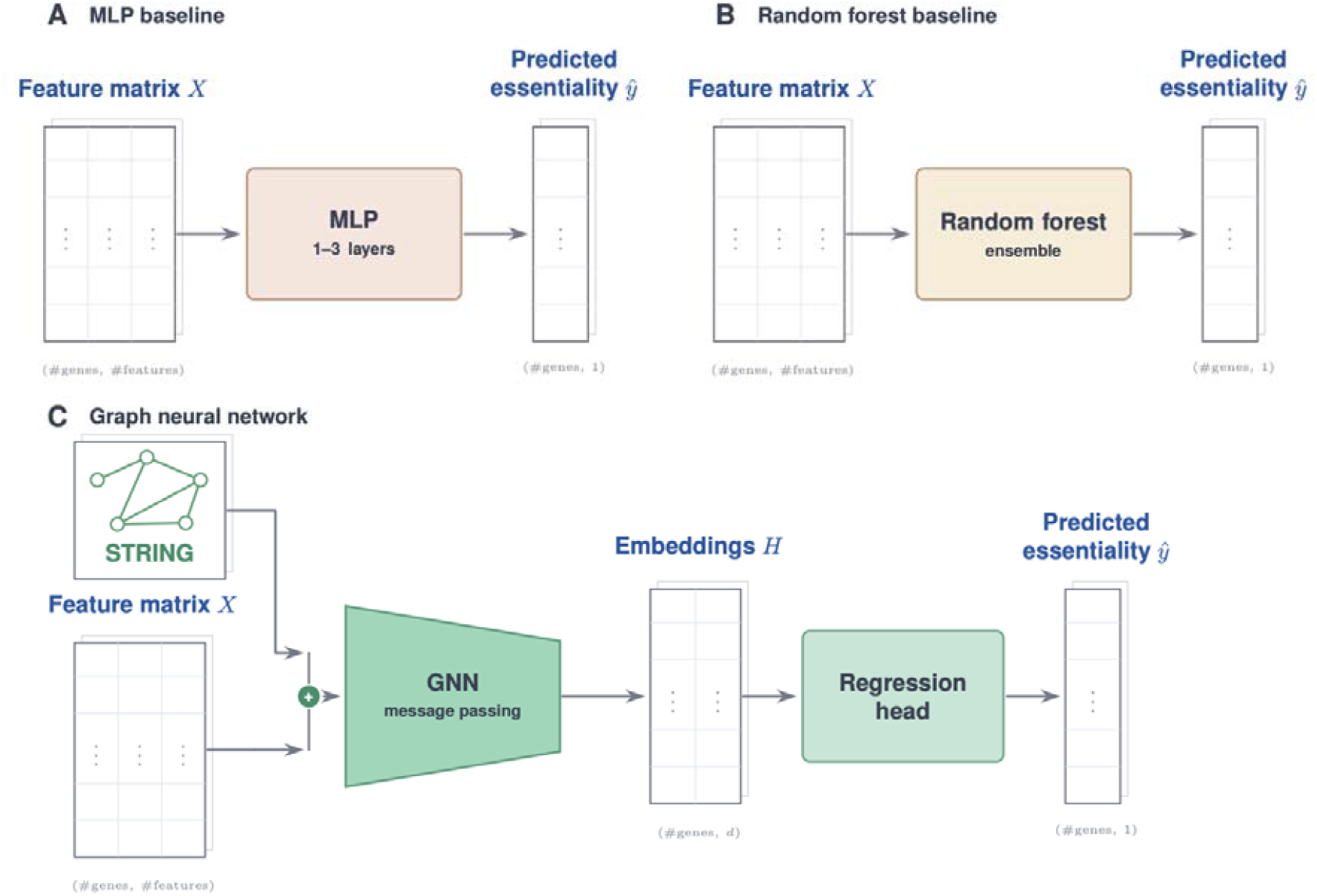
Overview of model architectures used for gene essentiality prediction. (a) Multilayer perceptron (MLP): operates on concatenated gene-level and cell-line–specific features to predict essentiality scores. (b) Random forest (RF): ensemble-based model using the same feature space without normalization. (c) Graph neural network (GNN): integrates node features with the gene–gene interaction graph (STRING) via message passing, producing gene embeddings that are mapped to essentiality scores through a regression head. All models are trained under identical gene-level splits and evaluation protocols.

A Random Forest regressor implemented in scikit-learn (Pedregosa et al., 2011) was trained on the same features as the MLP without normalization. Key hyperparameters (number of trees, maximum depth, minimum samples per split, feature subsampling) were optimized via Optuna (Akiba *et al*. 2019).

Graph Neural Networks (GNNs) operated on the homogeneous gene-gene interaction graph with node features as described above. We evaluated five message-passing architectures: GraphSAGE (Hamilton, Ying, and Leskovec 2017), Graph Attention Networks (GAT; Veličković *et al*. 2017), Graph Isomorphism Networks (GIN; Xu *et al*. 2018), Principal Neighbourhood Aggregation (PNA; Corso *et al*. 2020), and Graph Convolutional Networks (GCN; Kipf and Welling 2017). Architecture (GNN type), activation function, depth, hidden dimensionality, aggregation function, and regularization hyperparameters were selected via multi-objective Optuna optimization. All GNNs performed node-level prediction. Activation function, Batch Normalization, and dropout were applied to intermediate representations. The hyperparameter search space is summarized in **Table S1**.

We additionally evaluated a graph transformer architecture following the GraphGPS design (Rampášek et al., 2022), which combines local message passing with global self-attention across all nodes. GraphGPS was selected as a representative hybrid graph transformer because it integrates both neighborhood aggregation and graph-wide context, enabling a direct assessment of whether long-range dependencies provide additional predictive value beyond the local biological interactions captured by conventional GNNs. Each layer interleaves a local convolution (GraphSAGE or GIN) with a multi-head dense self-attention module followed by a feed-forward block. To inject graph-structural information that the permutation-invariant attention would otherwise lack, we used three positional encodings combined inside the input encoder: a learnable degree embedding (capturing hub connectivity), a Laplacian positional encoding (LapPE; Dwivedi and Bresson, 2021; Kreuzer et al., 2021) based on the top-k Laplacian eigenvectors (k 11 {8, 16, 32}) encoded via a DeepSets MLP over (|u|, u^2^) to address eigenvector sign ambiguity, and an optional random-walk positional encoding (RWPE; Dwivedi et al., 2022). LapPE was selected as the primary structural prior because protein– protein interaction networks exhibit modular organization that is well captured by the low-frequency spectrum of the graph Laplacian; degree adds local connectivity context essentially for free; RWPE provides local random-walk accessibility complementary to LapPE’s global view. The three encodings were summed with the projected node features before entering the GPS stack. All remaining hyperparameters (number of layers, hidden dimension, attention heads, learning rate, optimizer, scheduler, etc.) were tuned with Optuna over 100 trials per fold, using identical cross-validation splits, training procedure, and early-stopping criteria as the GNN baselines, so that the comparison reflects only the architectural difference.

### Graph Structure Controls and Feature Ablation

To isolate the contribution of biological network structure, we evaluated GNN models across three graph variants, while independently optimizing architecture and training hyperparameters for each condition to ensure a fair comparison. The graph variants included: (i) the biological STRING interaction network; (ii) a degree-preserving shuffled network; and (iii) a fully random graph with matched node and edge counts **(Figure S2)**. All three graphs share identical node sets and node features; only edge connectivity differs. Network-derived node features were computed once from the STRING network and held fixed across all graph conditions, allowing differences to be attributed specifically to the graph connectivity used during message passing rather than changes in node-level feature representations.

The degree-preserving shuffled network was generated using a double-edge swap procedure (Maslov and Sneppen, 2002), in which pairs of edges are (u, v) and (x, y) are rewired to (u, y) and (x, v) when valid. This process was iterated extensively (~10× the number of edges) to ensure sufficient randomization. This approach preserves the degree distribution while disrupting higher-order biological organization, including pathway structure and functional modules.

The fully random graph was constructed by uniformly sampling node pairs to match the number of edges in the STRING network, resulting in an Erdős–Rényi graph (Erdős and Rényi, 1959) with identical node and edge counts but no preserved degree structure or biological organization. Together, these controls enable separation of degree-related effects from higher-order biological connectivity.

For each null graph class (degree-preserving and fully random), a single realization was generated and used across all cross-validation folds to ensure that performance differences reflected model variability rather than variation in graph instantiation. While multiple realizations can better capture null variability, using a single instance combined with cross-validation is consistent with prior GNN studies in biological networks (e.g., Zitnik, Agrawal, and Leskovec 2018, Kulmanov and Hoehndorf 2020), where cross-validation provides a practical estimate of variability.

To further disentangle the contribution of graph structure from node-level information, we performed feature ablation within each graph condition. GNNs were trained under two feature settings: (a) the full feature set, including network-derived features (degree, closeness centrality, betweenness centrality, PageRank, and shortest-path distances), and (b) a reduced feature set excluding all such topological summaries. This design addresses two complementary questions:

First, whether biological network structure provides predictive signal beyond generic topology, assessed by comparing performance across STRING, shuffled, and random graphs within each feature setting. If the STRING network provides a meaningful biological signal, it should outperform shuffled and random graphs, particularly under the reduced feature setting where no pre-computed topological features are available.

Second, whether GNNs extract topological information through message passing or rely on pre-computed node features, assessed by comparing full and reduced feature sets within each graph. If meaningful structure is captured through connectivity, performance should be preserved without explicit network features; conversely, substantial degradation indicates reliance on node-level topological summaries. All graph variants share identical node features, differing only in edge connectivity.

### Training Procedure and Hyperparameter Optimization

Neural networks were implemented in PyTorch with PyTorch Geometric (Fey & Lenssen, 2019). Optimization used AdamW (Loshchilov and Hutter 2017) or RMSprop with either OneCycleLR or ReduceLROnPlateau scheduling. Training ran for up to 100 epochs with early stopping (patience = 8, minimum improvement threshold = 1e-3). Mini-batches corresponded to individual cell lines, such that all genes within a cell line were processed jointly in a single forward pass; this avoids the block-diagonal adjacency matrices that would arise from batching multiple cell lines.

Gradient accumulation steps were selected from {1, 2, 4} as a tunable hyperparameter. Mixed-precision training (AMP) was applied throughout.

Hyperparameters were optimized using Optuna (100 trials per tissue and model) in a multi-objective setting minimizing MSE and maximizing MCC. A single configuration was selected from the Pareto front based on minimum Euclidean distance to the ideal point after normalizing both objectives to [0, 1].

### Multi-Tissue Evaluation

All experiments were conducted independently for breast, lung, and large intestine tissues using identical pipelines. Hyperparameter optimization was performed separately per tissue to account for tissue-specific data distributions. Results are reported per tissue.

## Results

### Gene-level evaluation reveals consistent advantages of graph-based models over feature-only baselines

A key challenge in gene essentiality prediction is determining whether models can generalize to genes not previously observed during training. Conventional cell-line-level evaluation often places the same genes in both training and test sets across different cellular contexts, potentially inflating performance through gene-specific information leakage. Consistent with this, a simple gene-wise mean predictor achieved strong performance across all tissues under cell-line-level splits (**Table S2**), indicating that such evaluation substantially overestimates true generalization capability. To address this, all subsequent analyses use a strict gene-level split in which test genes are entirely unseen during training.

We evaluated all models independently across breast (n = 35 cell lines), lung (n = 43), and large intestine (n = 106) tissues, to test generalisation across biological context (**Table S3**). Under the stringent gene-level evaluation, the GraphSAGE-based GNN, identified as the best-performing architecture during hyperparameter optimization **(Table S4)**, consistently achieved the lowest MSE and highest MCC across all three tissues **(Figure 2; Table S5)**, demonstrating superior performance on both regression and classification objectives. The MLP achieved the second-lowest MSE but ranked last on MCC, indicating that while it captures continuous variation in essentiality, it is less effective at discriminating essential from non-essential genes. In contrast, the Random Forest showed higher MCC than the MLP but the highest MSE, suggesting that it retains some classification-relevant structure despite poorer regression accuracy. Only the GNN performs strongly across both objectives, making it the most balanced model under gene-level generalization. All pairwise differences between models were statistically significant (GNN vs MLP: p < 0.01; GNN vs RF and MLP vs RF: p < 0.001; **Table S6**).

**Figure 2.**
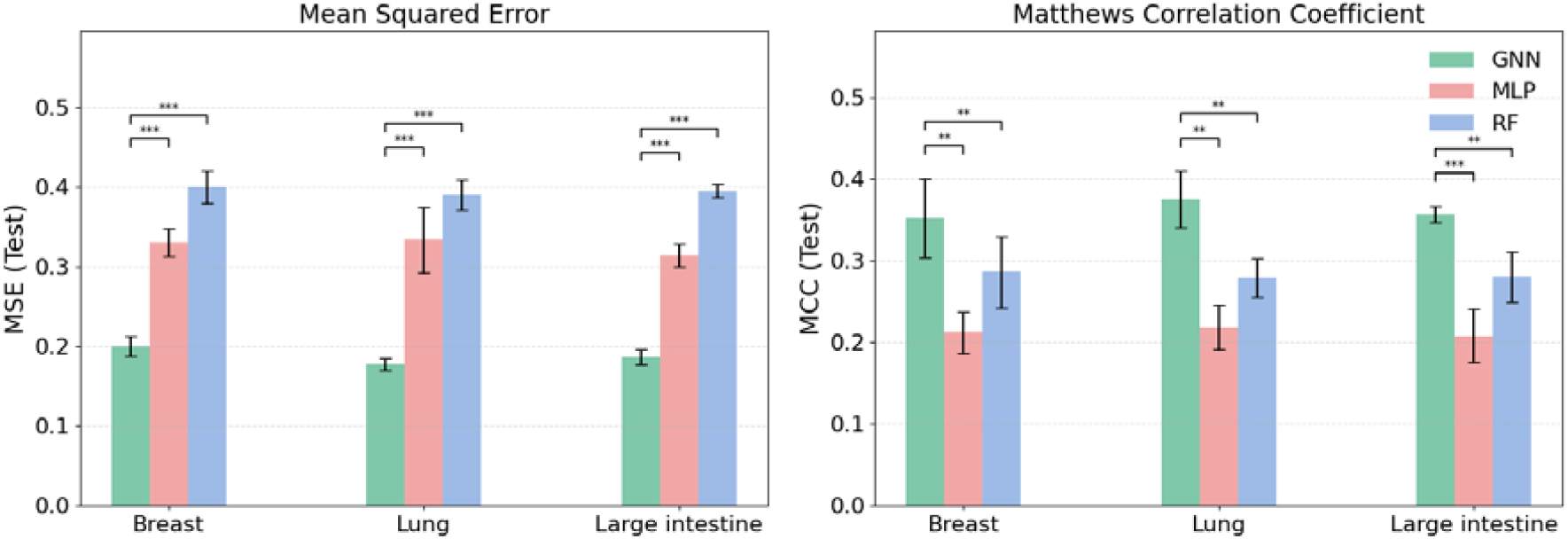
Cross-tissue comparison of graph-based and feature-only models under gene-level evaluation. Test-set performance of the graph neural network (GNN), multilayer perceptron (MLP), and random forest (RF) across breast, lung, and large intestine tissues. Performance is reported as mean squared error (MSE; left panel; lower is better) and Matthews correlation coefficient (MCC; right panel; higher is better). Bars represent mean performance across five cross-validation folds, with error bars indicating standard deviation. Statistical significance between model pairs is shown (**p < 0.01; ***p < 0.001; paired t-test).

This advantage was consistent despite differences in sample size, essentiality distributions, and tissue-specific dependency profiles (**Figure S3; Table S3**), indicating robustness to variation in dataset composition. The semi-supervised formulation allows the GNN to propagate information across the interaction network, providing a relational inductive bias that is not available to feature-only models operating independently per gene.

### Biological interaction structure, not generic topology, drives graph-based gains

To determine whether GNN improvements reflect biologically meaningful network organization or generic graph connectivity, we evaluated the GNN framework across three graph conditions: the widely used STRING functional protein association network, a degree-preserving shuffled network generated via double-edge swaps, and a fully random graph with matched node and edge counts. Architecture and training hyperparameters were independently optimized for each condition to ensure a fair comparison.

Disrupting biological network structure led to consistent performance degradation across tissues (**Figure 3; Table S7**). Relative to the STRING network, degree-preserving shuffled graphs increased MSE by 0.0239–0.0272 and reduced MCC by 0.0885–0.0962 (p = 0.014–0.038), while fully random graphs caused further deterioration (MSE increase: 0.0284–0.0407; MCC decrease: 0.0824–0.1372; p = 0.007–0.049; paired t-test across folds; **Table S8**).

**Figure 3.**
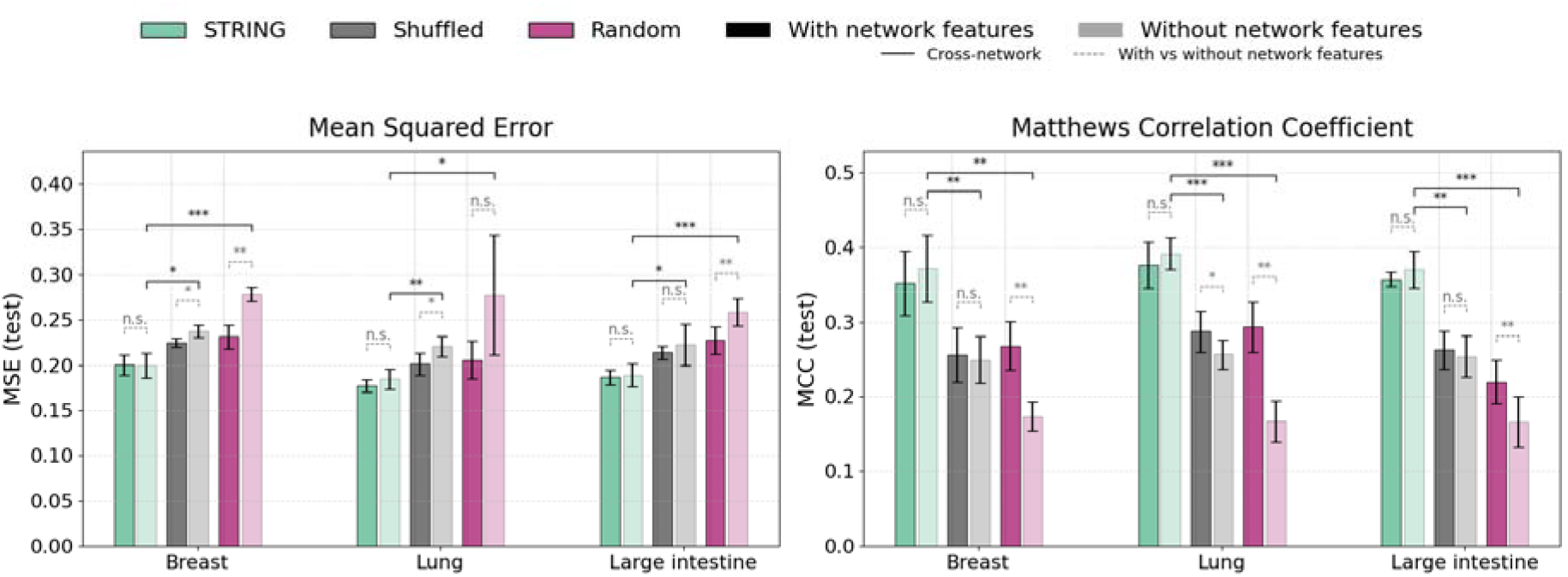
Disentangling biological network signal from generic topology in GNN performance. Test-set performance of the GNN under three graph conditions—STRING biological network, degree-preserving shuffled network, and fully random network—and two feature settings: with and without network-derived node features from STRING. Results are shown for breast, lung, and large intestine tissues. Left panels: mean squared error (MSE; lower is better). Right panels: Matthews correlation coefficient (MCC; higher is better). Bars represent mean ± standard deviation across five folds. Solid brackets indicate paired t-tests comparisons across graph types within the same feature condition (cross-network comparisons), while dashed brackets indicate comparisons between feature conditions within the same graph (feature ablation).

Critically, because the degree-preserving shuffled graph preserves node degree distribution while disrupting higher-order biological organization, the performance gap between STRING and the shuffled graph demonstrates that predictive gains cannot be attributed to degree effects or graph density alone. Instead, the progressive decline from biological to shuffled and random graphs indicates that graph-based gains arise from biologically structured gene–gene relationships, such as pathways and functional modules, rather than arbitrary connectivity.

### Message passing over STRING captures topological signal without requiring pre-computed network features

To further disentangle the source of graph-based gains, we performed a feature ablation within each graph condition, comparing GNN performance using the full feature set, including network-derived features such as degree, closeness centrality, betweenness centrality, and PageRank, against a reduced feature set excluding all such topological summaries (**Figure 3**, dashed brackets; **Table S9**). For the STRING network, removing network-derived features had negligible effect on performance across all tissues. Changes in MSE (Δ = −0.0004 to +0.0071), and MCC (Δ = +0.013–0.019) were small and not statistically significant, indicating that message passing over the biological graph is sufficient to capture topological signal without requiring explicit node-level encoding.

For the degree-preserving shuffled graph, the effect was moderate and inconsistent. MSE increased by 0.0089–0.0192 and MCC decreased by up to 0.0303, with significant degradation observed in three of six comparisons (Breast MSE, Lung MSE and MCC). This suggests that although biological organization is disrupted, residual degree-related structure can still be partially exploited through message passing.

In contrast, for the fully random graph, removing network-derived features led to substantial and mostly significant performance degradation, (MSE increase: 0.031-0.072; MCC decrease: 0.053-0.126), with all comparisons except Lung MSE being statistically significant. This indicates that when graph connectivity is uninformative, the model relies heavily on pre-computed topological node features to maintain predictive performance.

Taken together, these results demonstrate that topological signal is not inherent to message passing alone, but depends on biologically structured connectivity. In the STRING network, meaningful edge structure enables the GNN to learn topology implicitly, whereas under random connectivity, this signal must be externally supplied through node features.

### Local message passing outperforms combined local–global graph representation learning

We additionally evaluated a GraphGPS-style architecture, which combines local graph convolutions with global self-attention across all nodes. Across breast, lung, and large intestine tissues, despite the increased expressive capacity of transformer-based models, the standard GNN outperformed the GPS model on both MSE and MCC (**Figure 4**; **Table S10**). These differences were statistically significant in most cases (breast: p < 0.01; large intestine: p < 0.05), with the exception of lung MSE. These findings indicate that global attention does not improve performance over local message passing in this task, at least as implemented in GraphGPS. This is consistent with the modular organization of protein interaction networks, where functionally related genes are concentrated within local neighborhoods rather than distributed globally across the graph. Incorporating global attention may dilute this local signal or introduce noise, particularly under the gene-level generalization setting. More broadly, these results suggest that increased architectural complexity does not necessarily translate into improved performance in biologically structured graphs.

**Figure 4.**
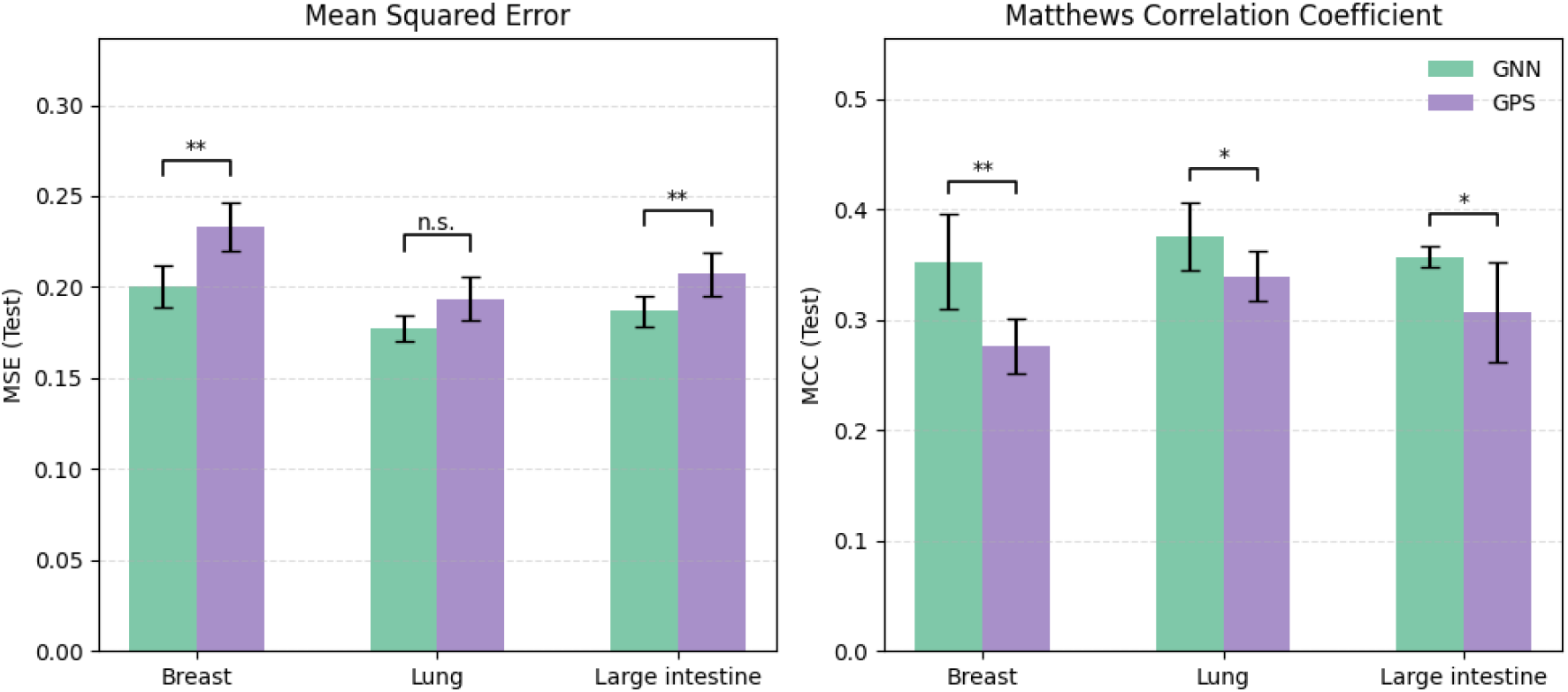
Comparison of local message passing and hybrid local–global attention architectures. Test-set performance of a standard graph neural network (GNN) and a GraphGPS-style model combining local message passing with global self-attention (GPS), evaluated across breast, lung, and large intestine tissues. Performance is reported as mean squared error (MSE; left panel; lower is better) and Matthews correlation coefficient (MCC; right panel; higher is better). Bars indicate mean ± standard deviation across five cross-validation folds. Statistical comparisons between models are shown (**p < 0.01; *p < 0.05; n.s., not significant; paired t-test).

### Limited within-gene variability constrains per-gene predictability

Per-gene Pearson and Spearman correlations between predicted and observed essentiality values across cell lines were centered near zero across all models and tissues, with broad and largely overlapping distributions (**Figures S4–S5**).

This pattern reflects a structural property of the task under gene-level splitting: for many genes, essentiality varies only modestly across cell lines, imposing a ceiling on the achievable within-gene correlation. As a result, model improvements are more apparent in global metrics, such as overall ranking and classification performance, rather than in fine-grained prediction of within-gene variability across contexts.

In this setting, the primary utility of predictive models lies in improved prioritization and discrimination of essential versus non-essential genes across unseen genomic contexts. Importantly, this limitation applies uniformly across all models, indicating that it reflects intrinsic data properties and limits of predictability, rather than model deficiencies.

## Discussion

Predicting gene essentiality across cellular contexts requires models to generalize beyond previously observed genes while accounting for complex, context-dependent biological dependencies. A central unresolved question is whether the apparent success of graph-based models reflects biologically meaningful interaction structure or simply generic properties of graph topology. Here, using a controlled evaluation framework combining gene-level splitting, topology-matched graph perturbations, and feature ablation, we show that graph neural networks (GNNs) outperform feature-only baselines under stringent generalization, but that these gains depend critically on biologically structured connectivity. Specifically, predictive performance degrades when biological networks are replaced with degree-preserving or random graphs, and network-derived node features become important only when graph structure is uninformative. In addition, global attention mechanisms do not improve performance over local message passing, and per-gene correlations remain uniformly low across models. Together, these findings indicate that graph-based gains arise from biologically meaningful local interaction structure, while also revealing fundamental limits of gene-level predictability.

A key implication is that evaluation design fundamentally shapes conclusions about model performance. Under conventional cell-line splits, models can exploit the relative stability of gene essentiality across contexts, leading to inflated performance that does not reflect true generalization. By adopting gene-level splits, we remove this shortcut and expose the intrinsic difficulty of predicting essentiality for unseen genes, consistent with observations in related domains such as drug response prediction (Haibe-Kains et al. 2013, Manica et al. 2019). These results underscore the importance of aligning evaluation frameworks with intended deployment scenarios.

The consistent advantage of GNNs over feature-only models highlights the importance of relational inductive bias in modeling gene function. Gene essentiality is not an intrinsic property of individual genes but emerges from their participation in functional systems, including protein complexes, pathways, and regulatory modules. Message passing incorporates this context by embedding each gene within its interaction neighborhood, effectively approximating the modular organization of cellular processes (Hart et al. 2015, Boyle, Li, and Pritchard 2017), and supporting the broader view that biological function is inherently networked rather than gene-centric.

Critically, our controlled perturbation experiments further demonstrate that these gains depend on biological organization rather than generic topology. Performance declines not only under fully random graphs, but also under degree-preserving rewiring, indicating that centrality alone cannot explain predictive performance, in contrast to previous observations (Jeong et al. 2001). Instead, the identity and specificity of interactions, capturing pathway co-membership, co-complex relationships, and functional dependencies, provide the dominant signal. This finding is consistent with systems-level models of cellular organization, in which function arises from structured, modular interactions rather than simple connectivity patterns (Barabási, Gulbahce, and Loscalzo 2011). More broadly, it underscores the importance of topology-matched controls when evaluating graph-based models in biology, as improvements cannot be assumed to reflect meaningful signals without explicitly ruling out structural confounders.

The feature ablation analysis provides additional mechanistic insight into how GNNs utilize topology. On a biologically meaningful network, performance is preserved without network-derived node features, indicating that message passing can recover topological information directly from biologically meaningful connectivity alone. In contrast, under random connectivity, the removal of these features leads to substantial performance degradation, demonstrating that node-level summaries such as degree or centrality act as proxies when the graph itself is uninformative. This dissociation clarifies a common ambiguity in graph-based learning: improvements are not driven by redundant encoding of topology in node features, but by the interaction between feature information and meaningful graph structure. Similar observations have been reported in protein function prediction, where graph connectivity enables models to capture relational signals that cannot be replicated by feature engineering alone (Zitnik, Agrawal, and Leskovec 2018, Kulmanov and Hoehndorf 2020).

The comparison with Graph Transformer architectures suggests that relevant biological signals operate at a local rather than global scale. Despite their greater expressive capacity, models incorporating global self-attention do not outperform local message passing, consistent with the organization of protein interaction networks into localized functional modules (Boyle, Li, and Pritchard 2017). Global attention may introduce irrelevant long-range information that weakens locally informative signals, particularly under gene-level generalization where spurious correlations are harder to exploit. These findings indicate that model design should reflect the scale of biological organization, and that local message passing remains well-suited for node-level prediction tasks in dense biological networks.

Finally, uniformly low per-gene correlations highlight a fundamental limitation of the prediction task. Under gene-level evaluation, many genes exhibit limited variability across cell lines, constraining achievable within-gene correlation. This limitation is intrinsic to current CRISPR screening data and applies uniformly across model classes, indicating that it cannot be overcome by architectural improvements alone. Instead, progress in predicting context-specific variation is likely to depend on richer representations of gene–context interactions, including higher-resolution molecular features, larger and more diverse datasets, or perturbation designs that more directly probe conditional dependencies. In practical terms, this suggests that current models are better suited for global prioritization of essential genes rather than precise prediction of context-dependent variability.

Several limitations warrant consideration. The analysis is restricted to single-gene perturbations and may not generalize to combinatorial or synthetic lethal interactions, where network effects could be more pronounced. The STRING interaction network, while high-confidence, is incomplete and biased toward well-studied genes, potentially limiting generalizability. In addition, the interaction network was treated as static across cellular contexts, whereas context-specific interaction rewiring may further influence dependency relationships. Finally, only a single realization of each null graph was evaluated; although cross-validation provides a measure of variability, future work could strengthen conclusions by averaging across multiple randomizations.

In summary, this study demonstrates that biological interaction networks provide genuine and reproducible predictive signals for gene essentiality, and that this signal arises from structured, functionally meaningful connectivity rather than generic graph properties. By combining rigorous evaluation design with controlled perturbations of graph structure, we provide a principled framework for separating biologically meaningful connectivity from generic graph topology in graph-based gene essentiality prediction. These findings suggest that future progress in gene essentiality prediction will depend not only on model development, but also on improved network representations, richer context-specific data, and evaluation strategies that faithfully reflect real-world generalization challenges.

## Data and code availability

All datasets used in this study are publicly available. Gene essentiality and molecular profiling data were obtained from the DepMap Public 24Q2 release (Broad Institute, 2024), and protein–protein interaction data were obtained from STRING v12.0.

Code used for data processing, model training, and analysis is available on github at https://github.com/Whole-Cell-Signalling/gnn-sko-essentiality.

## Funding

The work was supported through core funding by the European Molecular Biology Laboratory (SR, EP). This work was funded by AstraZeneca through the EAZPOD programme (EP/SR).

## Acknowledgements

EMBL IT Support is acknowledged for the provision of computer and data storage servers. We also thank Dr. Heiko Horn and Dr. Dom Kirkham for helpful discussions and feedback.

## Author Contributions

S.R.: Conceptualization, Methodology, Software, Validation, Formal analysis, Investigation, Data curation, Visualization, Writing - original draft, Writing - review & editing. A.A. and S.B.: Methodology, Supervision, Writing - review & editing. M.M.: Supervision, Writing - review & editing. E.M.: Methodology, Supervision, Writing - review & editing. E.P.: Conceptualization, Methodology, Supervision, Project administration, Resources, Writing - review & editing.

## Conflict of Interest

This work was funded by AstraZeneca. SB, MM and EB are employees of AstraZeneca and EP of GlaxoSmithKline.

## Notes

### Competing Interest Statement

Stephen Bonner, Avid Afzal, Edward Morrissey and Marta Milo are employed by AstraZeneca and Evangelia Petsalaki is employed by GlaxoSmithKline.

### Summary of Updates

I forgot to add the name of one of the authors Avid Afzal in the list of the authors so I have now added them

