## Supplementary Information for "Biological Network Organization, Not Generic Graph Topology, Drives Graph-Based Gene Essentiality Prediction"

### Supplementary Materials

#### Table of Contents:

##### Supplementary Text

Supplementary Text S1. Cell-line split baseline reveals gene identity dominance

##### Supplementary Figures

Figure S1. Comparison of evaluation strategies: cell-line split vs gene-level split.

Figure S2. Schematic representation of graph structure controls used in this study.

Figure S3. Distribution of gene essentiality scores across tissues.

Figure S4. Distribution of per-gene Spearman correlations across models, tissues, and gene groups.

Figure S5. Distribution of per-gene Pearson correlations across models, tissues, and gene groups.

### Supplementary Tables

Table S1. Hyperparameter search space.

Table S2. Performance of gene-wise mean predictor under 5-fold CV cell-line split

Table S3. Full performance metrics for main model comparison (STRING network, full features).

Table S4. Statistical comparison of model performance on the Breast, Lung, and Large Intestine.

Table S5. Summary statistics of gene essentiality data across tissues.

Table S6. Full GNN performance under graph-structure and feature-ablation conditions.

Table S7. Impact of graph topology on GNN performance.

Table S8. Effect of network structure on GNN performance across tissues.

Table S9. Comparison of GNN and Graph Transformer (GPS) architectures.

Table S10.Summary of hyperparameter configurations across tissues and cross-validation folds.

#### Supplementary Text

***Text S1: Cell-line split baseline reveals gene identity dominance***

Under a cell-line split, we implemented a simple gene-wise mean predictor, where each gene's average essentiality across training cell lines was used as a constant prediction for unseen cell lines. This baseline does not use any input features, graph structure, or cell-line-specific information, relying solely on target values observed during training. Despite its simplicity, this approach achieved remarkably high performance across both regression and classification metrics (Table S1), with test Pearson correlation exceeding 0.88 and MCC above 0.70. These results indicate that a large fraction of the variance in gene essentiality can be explained by gene-specific effects that are consistent across cell lines. This finding highlights a critical limitation of cell-line split evaluation: models can achieve strong performance by effectively memorizing gene identity rather than learning biologically meaningful patterns or context-dependent dependencies. As a result, this setup does not adequately test generalization to unseen genes. To address this issue, we adopt a gene-level split throughout the main analysis, ensuring that all test genes are entirely unseen during training. This evaluation framework prevents reliance on gene-specific averages and more accurately reflects the ability of models to generalize across genes. All reported test-set metrics and per-gene statistics are computed exclusively on held-out test genes for each fold. Values are aggregated across folds after ensuring no gene appears in both train and test sets.

#### Supplementary Figures

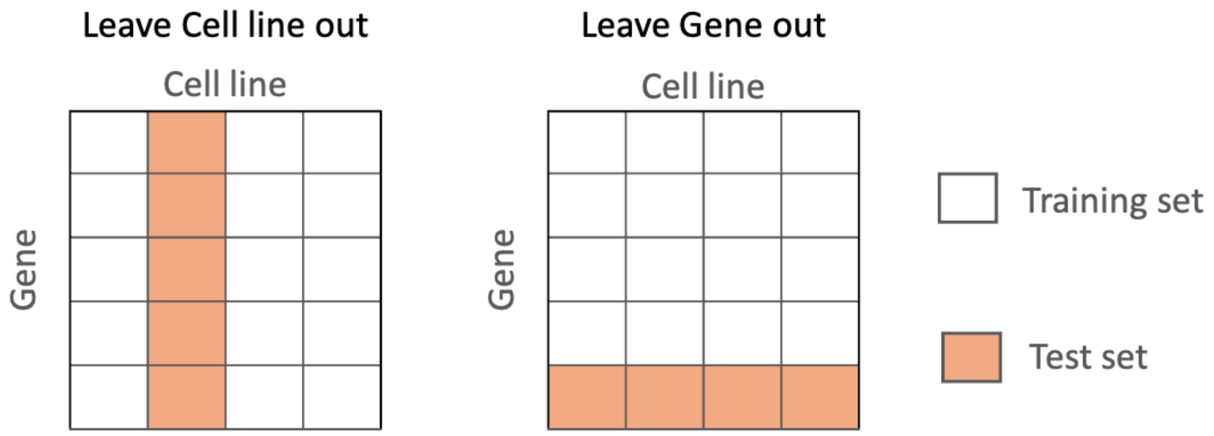


***Figure S1*. *Comparison of evaluation strategies: cell-line split vs gene-level split.*** *Illustration of two data splitting approaches. In the cell-line split (left), all genes appear in both training and test sets across different cell lines, allowing models to exploit gene-specific averages and leading to inflated performance estimates. In the gene-level split (right), entire genes are held out from training, requiring models to generalize to unseen genes. All results in this study are based on the gene-level split, providing a more stringent and biologically meaningful evaluation of model generalization.*

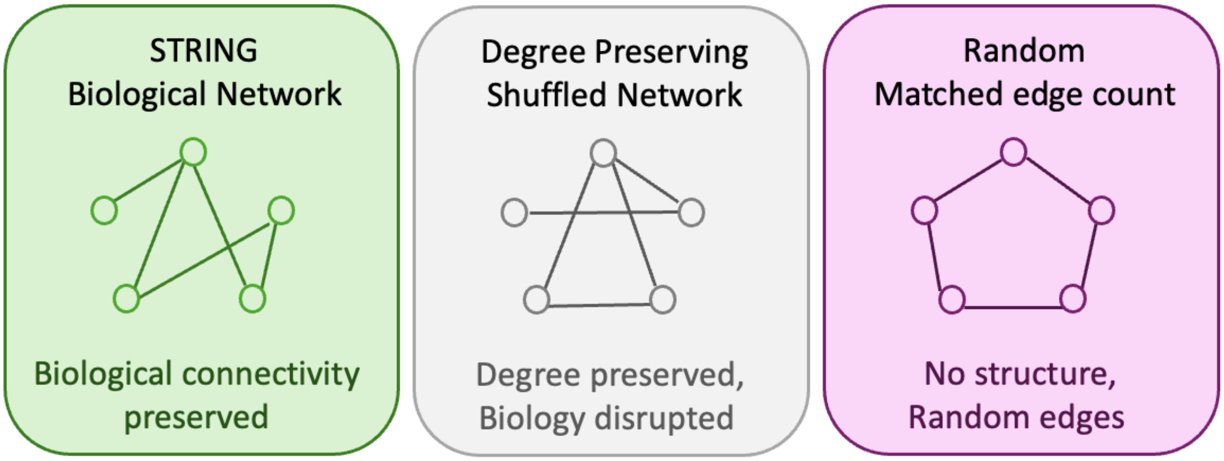


***Figure S2**. **Schematic representation of graph structure controls used in this study.*** *Three graph conditions used to isolate the contribution of biological structure in gene essentiality prediction: (i) the STRING biological interaction network, preserving functional relationships between genes; (ii) a degree-preserving shuffled network, in which edges are rewired while maintaining node degree, disrupting biological organization but preserving connectivity statistics; and (iii) a fully random network with matched node and edge counts, lacking both biological structure and meaningful topology. This controlled design enables separation of biologically meaningful signals from generic graph properties.*

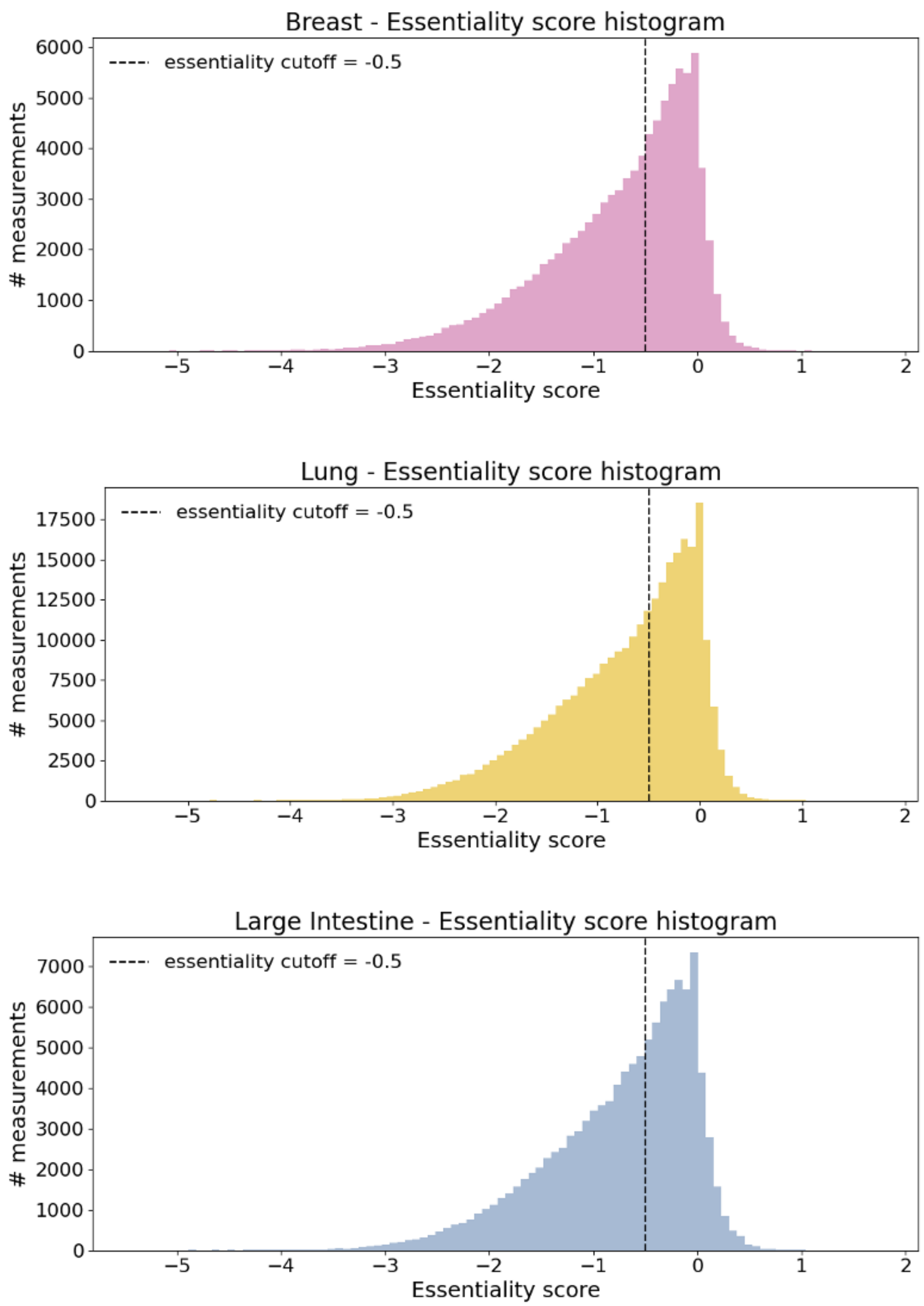


***Figure S3**. **Distribution of gene essentiality scores across tissues.** Histograms of gene essentiality scores for breast, lung, and large intestine tissues, aggregated over all (gene, cell line) pairs. The dashed vertical line indicates the threshold used to define essential genes (≤ −0.5). Across all tissues, score distributions are left-skewed, with a higher density of values near zero and a long tail toward strongly negative values, reflecting varying degrees of gene dependency. Differences in distribution shape and sample size across tissues highlight heterogeneity in essentiality profiles used for model evaluation. All genes are the same across the tissues.*

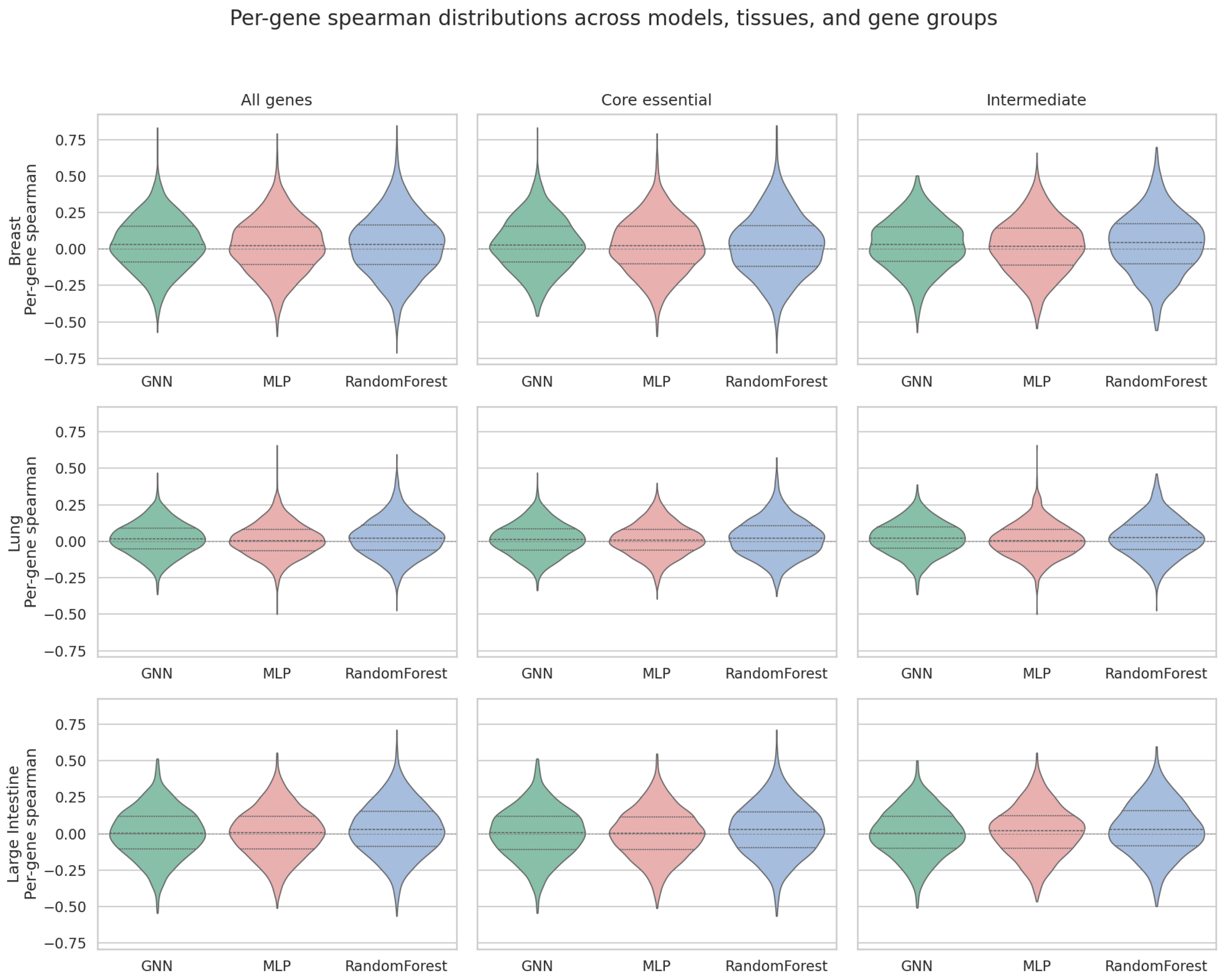


***Figure S4*. *Distribution of per-gene Spearman correlations across models, tissues, and gene groups.*** *Per-gene Spearman rank correlations were computed across cell lines for each held-out test gene, evaluating the ability of models to preserve relative ordering of essentiality values. Distributions are shown for GNN, MLP, and Random Forest models across tissues and gene groups. Consistent with Pearson results (Figure S2), Spearman correlations are centered near zero across all conditions, indicating that models struggle to recover gene-specific ranking patterns across cell lines. This suggests that predictive performance arises primarily from global signal rather than accurate modeling of fine-grained, gene-specific context-dependent differences. Similar behavior across model classes supports the conclusion that this limitation is driven by data characteristics rather than modeling approach.*

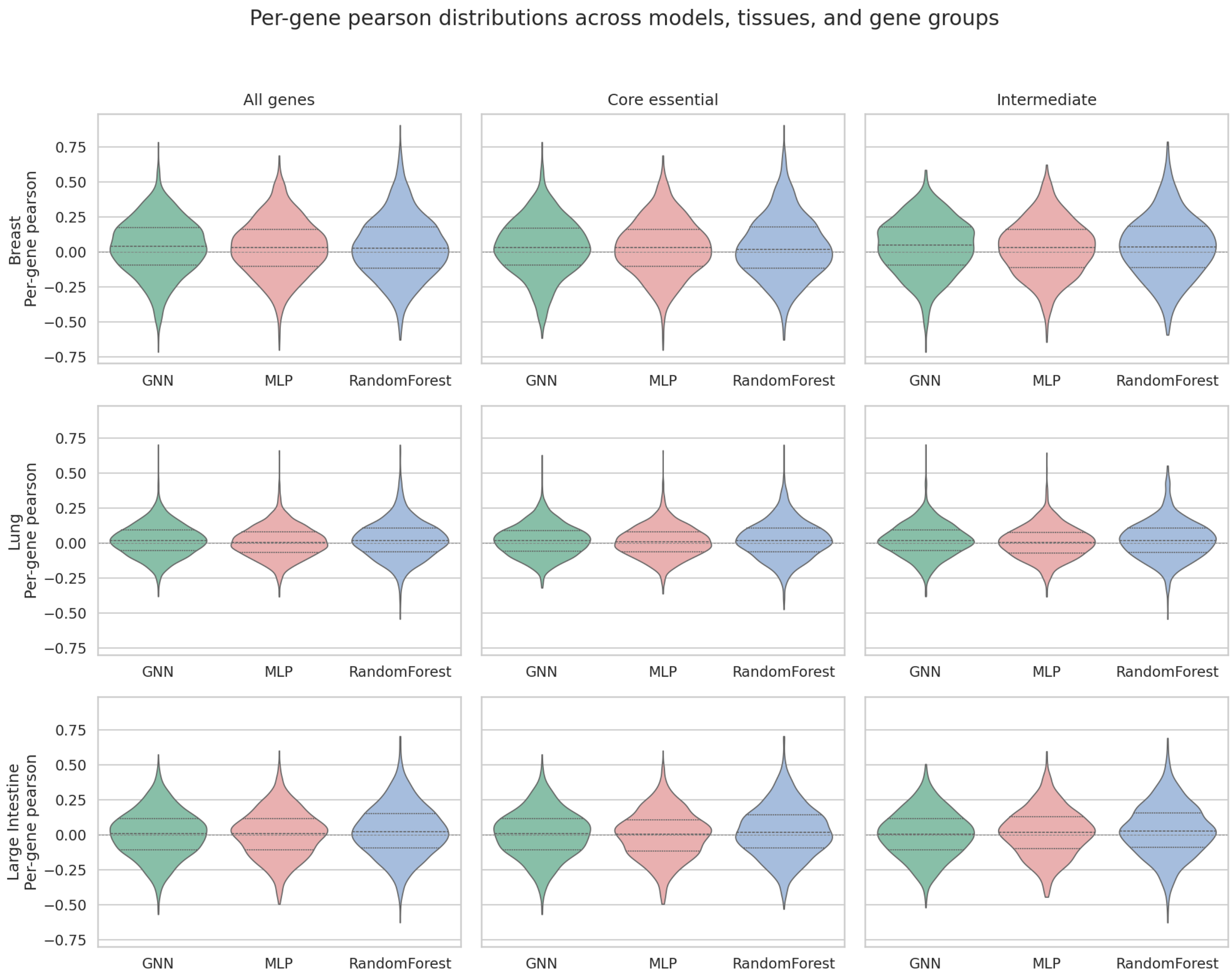


***Figure S5. Distribution of per-gene Pearson correlations across models, tissues, and gene groups.*** *Per-gene Pearson correlation coefficients were computed across cell lines for each held-out test gene under gene-split evaluation. Distributions are shown for three models (GNN, MLP, Random Forest) across tissues (Breast, Lung, Large intestine) and gene groups (All genes, Core essential, Intermediate). Across all tissues and gene groups, distributions are centered near zero, indicating limited ability of all models to capture gene-specific variability across cell lines. While global predictive performance (e.g., ranking and classification) is strong, these results suggest that improvements are driven primarily by capturing shared patterns across genes rather than accurately modeling within-gene context-dependent variation. No substantial differences are observed between models, further indicating that this limitation reflects intrinsic properties of the data rather than model architecture.*

### Supplementary Tables

***Table S1*. *Hyperparameter search space.*** *Hyperparameters optimized using Optuna for GNN, GPS, MLP, Ridge Regression, and Random Forest models. Continuous parameters were sampled on a log-uniform scale where indicated. Fixed parameters are reported as single values. For PNA-based GNNs, aggregators and scalers were fixed to the listed sets.*

| Model | Hyperparameter | Search space / value |
|---|---|---|
| **GNN / GPS / MLP** | Epochs | 100 |
| **GNN / GPS / MLP** | Optimizer | RMSprop, AdamW |
| **GNN / GPS / MLP** | Scheduler | OneCycleLR, ReduceLROnPlateau |
| **GNN / GPS / MLP** | Learning rate | log-uniform, 1e-4 to 3e-3 |
| **GNN / GPS / MLP** | Weight decay | log-uniform, 1e-6 to 1e-3 |
| **GNN / GPS / MLP** | Dropout probability p | 0, 0.1, 0.2 |
| **GNN / GPS / MLP** | Activation function | ReLU, PReLU |
| **GNN / GPS / MLP** | Accumulation steps | 1, 2, 4 |
| **GNN / GPS / MLP** | RMSprop alpha | 0.8 |
| **GNN / GPS / MLP** | RMSprop momentum | 0.6 |
| **GNN / GPS / MLP** | RMSprop eps | 1e-6 |
| **GNN** | GNN type | PNA, GraphSAGE, GIN, GCN, GAT |
| **GNN** | GNN hidden dimension | 64, 128, 256 |
| **GNN** | Output embedding dimension | 32, 64, 128 |
| **GNN** | Number of GNN layers | 2, 3, 4 |
| **GNN** | Regression head layers | 1, 2, 3 |
| **GNN** | PNA aggregators | mean, max |
| **GNN** | PNA scalers | identity, amplification, attenuation |
| **GPS** | Local convolution type | GraphSAGE, GIN |
| **GPS** | Hidden dimension | 32, 64, 128 |
| **GPS** | Output embedding dimension | 32, 64, 128 |
| **GPS** | Number of GPS layers | 1, 2, 3 |
| **GPS** | Regression head layers | 1, 2 |
| **GPS** | Attention heads | 1, 2 |
| **GPS** | Attention type | multihead |
| **GPS** | Laplacian PE dimension lap_pe_k | 8, 16, 32 |
| **GPS** | Use random-walk PE | True, False |
| **GPS** | Random-walk PE dimension rwpe_k | 8, 16 if RWPE enabled; otherwise 16 |
| **GPS** | Maximum degree embedding | 512 |
| **MLP** | Regression head layers | integer, 1 to 5 |

| | | |
|---|---|---|
| **Random Forest** | Number of trees | 100, 200 |
| **Random Forest** | Maximum depth | 8, 10, 12 |
| **Random Forest** | Minimum samples split | 5, 10, 20 |
| **Random Forest** | Minimum samples leaf | 2, 5 |
| **Random Forest** | Maximum features | sqrt, 0.3 |

***Table S2. Performance of gene-wise mean predictor under 5-fold CV cell-line split***

| Metric | Train | Test |
|---|---|---|
| **RMSE** | 0.3177 ± 0.0094 | 0.3295 ± 0.0350 |
| **Pearson** | 0.8984 ± 0.0053 | 0.8901 ± 0.0214 |
| **Spearman** | 0.8881 ± 0.0039 | 0.8790 ± 0.0157 |
| **BalancedAcc** | 0.6735 ± 0.0029 | 0.6664 ± 0.0176 |
| **F1_macro** | 0.6964 ± 0.0027 | 0.6873 ± 0.0179 |
| **F1_weighted** | 0.7850 ± 0.0035 | 0.7769 ± 0.0186 |
| **MCC** | 0.6296 ± 0.0050 | 0.6160 ± 0.0235 |

***Table S3. Summary statistics of gene essentiality data across tissues.*** *Descriptive statistics of Chronos gene essentiality scores for breast, lung, and large intestine tissues after preprocessing. Each entry corresponds to all (gene, cell line) pairs within a tissue. Reported values include the number of cell lines, genes, and total samples, along with distributional properties of essentiality scores (mean, standard deviation, minimum, and maximum). Genes are classified as essential or non-essential using a threshold of −0.5, and counts for each category are reported. Differences in sample size and score distributions across tissues highlight variability in dataset composition used for model evaluation.*

| **Metric** | **Breast** | **Lung** | **Large Intestine** |
|---|---|---|---|
| **Number of cell lines** | 35 | 106 | 43 |
| **Number of genes** | 2,741 | 2,741 | 2,741 |
| **Number of samples** | 95,935 | 290,546 | 117,863 |
| **mean** | -0.749 | -0.736 | -0.743 |
| **std** | 0.724 | 0.698 | 0.709 |
| **min** | -5.657 | -5.218 | -5.483 |
| **max** | 1.609 | 1.969 | 1.936 |
| **Number of essential genes(≤ -0.5)** | 51,659 | 156,723 | 64,099 |
| **Number of non_essential (> -0.5)** | 44,276 | 133,823 | 53,764 |

***Table S4. Summary of hyperparameter configurations across tissues and cross-validation folds.*** *Hyperparameter optimization was performed using Optuna over predefined search spaces. The table reports the range of selected values across all tissues (Breast, Lung, Large Intestine) and folds. GNN models consistently selected GraphSAGE architectures with moderate depth (2–4 layers) and hidden dimensions (128–256), while optimization parameters (learning rate, weight decay) varied across folds. MLP configurations were highly stable, converging to fixed architectures (3 layers, 64 hidden units). Random Forest models showed limited variability due to discrete search spaces. GPS models introduced additional structural encoding parameters but did not exhibit consistent performance advantages. These results indicate that GNN performance gains are robust and not driven by hyperparameter selection.*

| Model | Parameter | Range observed | Notes |
|---|---|---|---|
| **All (NN models)** | Optimizer | RMSprop (dominant), AdamW (rare) | RMSprop consistently selected |
| | Scheduler | ReduceLROnPlateau, OneCycleLR | Plateau most frequent |
| | Learning rate | 4.4e-4 → 2.7e-3 | Moderate variation |
| | Weight decay | 2e-5 → 6.5e-4 | Regularization varies |
| | Dropout (p) | 0.0 – 0.2 | Low dropout preferred |
| | Epochs | 50 – 100 | Stable |
| **GNN (STRING)** | GNN type | GraphSAGE | **Consistent across all tissues** |
| | Hidden dim (cl) | 128 – 256 | Larger embeddings preferred |
| | Layers (cl branch) | 2 – 4 | Moderate depth |
| | Regression layers | 1 – 3 | Shallow head |
| | Output dim | 32 – 128 | Flexible |
| | Activation | ReLU, PReLU | Minor variation |
| **GPS** | Local conv | SAGE, GIN | Hybrid local message passing |
| | Hidden dim | 32 – 128 | Smaller than GNN |
| | Layers | 1 – 3 | Shallower than GNN |
| | Attention heads | 1 – 2 | Low-head regime |
| | Laplacian PE (k) | 8 – 32 | Structural encoding |
| | RWPE | True / False | Optional |
| | RWPE k | 8 – 16 | When enabled |
| | Max degree embed | 512 | Fixed |
| **MLP** | Hidden dim | 64 | **Fully stable across tissues** |

| | | | |
|---|---|---|---|
| | Layers | 3 | **Fully stable** |
| | Learning rate | ~5e-4 – 1e-3 | Narrow range |
| | Dropout | 0.0 – 0.2 | Minor variation |
| **Random Forest** | n_estimators | 100 – 200 | Moderate ensemble size |
| | Max depth | 8 – 12 | Controlled tree depth |
| | Min samples split | 5 – 20 | Regularization |
| | Min samples leaf | 2 – 5 | Regularization |
| | Max features | sqrt, 0.3 | Feature subsampling |

***Table S5**. **Full performance metrics for main model comparison (STRING network, full features)**. Regression and classification performance of GNN, MLP, and Random Forest models using the STRING interaction network and full feature set. Results are averaged over five gene-level cross-validation folds and reported as mean ± standard deviation. Metrics include RMSE, Pearson and Spearman correlation, balanced accuracy, F1 scores (macro and weighted), and Matthews correlation coefficient (MCC), evaluated on both training and test sets.*

| **Tissue: Breast** | | | | | | |
|---|---|---|---|---|---|---|
| **Metric** | **GNN Train** | **GNN Test** | **MLP Train** | **MLP Test** | **RF Train** | **RF Test** |
| **MSE** | 0.0777 ± 0.0139 | 0.2003 ± 0.0112 | 0.1099 ± 0.0041 | 0.3305 ± 0.0158 | 0.1563 ± 0.0064 | 0.4005 ± 0.0178 |
| **Pearson** | 0.8302 ± 0.0316 | 0.5047 ± 0.0528 | 0.7665 ± 0.0094 | 0.3410 ± 0.0312 | 0.8455 ± 0.0076 | 0.4886 ± 0.0389 |
| **Spearman** | 0.8207 ± 0.0382 | 0.4867 ± 0.0510 | 0.7425 ± 0.0086 | 0.3194 ± 0.0306 | 0.8241 ± 0.0074 | 0.4575 ± 0.0327 |
| **Balanced Accuracy** | 0.8245 ± 0.0266 | 0.6734 ± 0.0231 | 0.7694 ± 0.0089 | 0.6022 ± 0.0109 | 0.8095 ± 0.0086 | 0.6326 ± 0.0183 |
| **F1 (macro)** | 0.8248 ± 0.0257 | 0.6727 ± 0.0241 | 0.7712 ± 0.0089 | 0.5995 ± 0.0113 | 0.8120 ± 0.0082 | 0.6265 ± 0.0222 |
| **F1 (weighted)** | 0.8262 ± 0.0250 | 0.6764 ± 0.0246 | 0.7739 ± 0.0085 | 0.6055 ± 0.0117 | 0.8143 ± 0.0077 | 0.6335 ± 0.0250 |
| **MCC** | 0.6509 ± 0.0495 | 0.3524 ± 0.0430 | 0.5496 ± 0.0143 | 0.2120 ± 0.0222 | 0.6323 ± 0.0139 | 0.2864 ± 0.0390 |
| **Tissue: Lung** | | | | | | |
| **Metric** | **GNN Train** | **GNN Test** | **MLP Train** | **MLP Test** | **RF Train** | **RF Test** |

| | | | | | | |
|---|---|---|---|---|---|---|
| **RMSE** | 0.0606 ± 0.0091 | 0.1774 ± 0.0070 | 0.0752 ± 0.0085 | 0.3340 ± 0.0367 | 0.1526 ± 0.0064 | 0.3902 ± 0.0174 |
| **Pearson** | 0.8637 ± 0.0226 | 0.5343 ± 0.0243 | 0.8313 ± 0.0175 | 0.3157 ± 0.0433 | 0.8345 ± 0.0100 | 0.4755 ± 0.0192 |
| **Spearman** | 0.8512 ± 0.0249 | 0.5105 ± 0.0321 | 0.8038 ± 0.0219 | 0.3027 ± 0.0366 | 0.8082 ± 0.0117 | 0.4393 ± 0.0255 |
| **Balanced Accuracy** | 0.8477 ± 0.0143 | 0.6879 ± 0.0155 | 0.8197 ± 0.0107 | 0.6086 ± 0.0128 | 0.8018 ± 0.0104 | 0.6328 ± 0.0113 |
| **F1 (macro)** | 0.8462 ± 0.0143 | 0.6871 ± 0.0151 | 0.8190 ± 0.0116 | 0.6079 ± 0.0138 | 0.8041 ± 0.0104 | 0.6291 ± 0.0156 |
| **F1 (weighted)** | 0.8469 ± 0.0143 | 0.6891 ± 0.0152 | 0.8200 ± 0.0119 | 0.6116 ± 0.0141 | 0.8063 ± 0.0101 | 0.6350 ± 0.0185 |
| **MCC** | 0.6937 ± 0.0286 | 0.3756 ± 0.0307 | 0.6390 ± 0.0223 | 0.2188 ± 0.0245 | 0.6149 ± 0.0195 | 0.2793 ± 0.0211 |
| **Tissue: Large Intestine** | | | | | | |
| **Metric** | **GNN Train** | **GNN Test** | **MLP Train** | **MLP Test** | **RF Train** | **RF Test** |
| **MSE** | 0.0807 ± 0.0072 | 0.1867 ± 0.0082 | 0.0930 ± 0.0064 | 0.3144 ± 0.0128 | 0.1546 ± 0.0023 | 0.3957 ± 0.0073 |
| **Pearson** | 0.8238 ± 0.0172 | 0.5101 ± 0.0099 | 0.7921 ± 0.0162 | 0.3249 ± 0.0225 | 0.8396 ± 0.0041 | 0.4791 ± 0.0194 |
| **Spearman** | 0.8130 ± 0.0173 | 0.4922 ± 0.0100 | 0.7679 ± 0.0182 | 0.3126 ± 0.0296 | 0.8137 ± 0.0042 | 0.4440 ± 0.0177 |
| **Balanced Accuracy** | 0.8224 ± 0.0118 | 0.6777 ± 0.0045 | 0.7821 ± 0.0154 | 0.6003 ± 0.0133 | 0.8068 ± 0.0086 | 0.6327 ± 0.0136 |
| **F1 (macro)** | 0.8216 ± 0.0113 | 0.6778 ± 0.0047 | 0.7840 ± 0.0153 | 0.5982 ± 0.0140 | 0.8092 ± 0.0084 | 0.6298 ± 0.0148 |
| **F1 (weighted)** | 0.8229 ± 0.0111 | 0.6808 ± 0.0055 | 0.7868 ± 0.0148 | 0.6049 ± 0.0156 | 0.8117 ± 0.0082 | 0.6369 ± 0.0148 |
| **MCC** | 0.6439 ± 0.0228 | 0.3570 ± 0.0092 | 0.5739 ± 0.0264 | 0.2082 ± 0.0289 | 0.6241 ± 0.0140 | 0.2807 ± 0.0274 |

***Table S6*. *Statistical comparison of model performance on the Breast, Lung, and Large Intestine.*** *Pairwise comparisons between models were performed using a two-sided paired Student's t-test (scipy.stats.ttest_rel) on per-fold test-set metrics across five cross-validation folds. Pairing was performed by fold identity, ensuring that each comparison reflects performance differences on identical test gene sets. Δ denotes the difference in mean performance (A − B), providing an estimate of effect size in addition to statistical significance.. Significance levels: * p < 0.05, ** p < 0.01, *** p < 0.001. No multiple-testing correction was applied, as comparisons were predefined and hypothesis-driven; most significant results remain robust at p < 0.01.*

| **Tissue: Breast** | | | | |
|---|---|---|---|---|
| | **Statistic** | **GNN vs MLP** | **GNN vs RF** | **MLP vs RF** |
| **MSE** | Mean A | 0.2003 | 0.2003 | 0.3305 |
| | Mean B | 0.3305 | 0.4005 | 0.4005 |
| | Δ (A–B) | -0.1302 | -0.2002 | -0.0700 |
| | t-stat | -12.63 | -17.47 | -6.38 |
| | p-value | 0.0002 | 0.0001 | 0.0031 |
| | Significance | *** | *** | ** |
| **MCC** | Mean A | 0.3524 | 0.3524 | 0.2120 |
| | Mean B | 0.2120 | 0.2864 | 0.2864 |
| | Δ (A–B) | +0.1404 | +0.0660 | -0.0744 |
| | t-stat | 7.76 | 6.16 | -4.17 |
| | p-value | 0.0015 | 0.0035 | 0.0141 |
| | Significance | ** | ** | * |
| **Tissue: Lung** | | | | |
| **MSE** | Mean A | 0.1774 | 0.1774 | 0.3340 |
| | Mean B | 0.3340 | 0.3902 | 0.3902 |
| | Δ (A–B) | -0.1567 | -0.2129 | -0.0562 |
| | t-stat | -9.49 | -25.60 | -3.71 |
| | p-value | 0.0007 | <0.0001 | 0.0206 |
| | Significance | *** | *** | * |
| **MCC** | Mean A | 0.3756 | 0.3756 | 0.2188 |

| | | | | |
|---|---|---|---|---|
| | Mean B | 0.2188 | 0.2793 | 0.2793 |
| | Δ (A–B) | +0.1568 | +0.0963 | -0.0605 |
| | t-stat | 6.71 | 4.71 | -5.56 |
| | p-value | 0.0026 | 0.0092 | 0.0051 |
| | Significance | ** | ** | ** |
| **Tissue: Large Intestine** | | | | |
| **MSE** | Mean A | 0.1867 | 0.1867 | 0.3144 |
| | Mean B | 0.3144 | 0.3957 | 0.3957 |
| | Δ (A–B) | -0.1277 | -0.2089 | -0.0812 |
| | t-stat | -15.20 | -47.42 | -10.21 |
| | p-value | 0.0001 | <0.0001 | 0.0005 |
| | Significance | *** | *** | *** |
| **MCC** | Mean A | 0.3570 | 0.3570 | 0.2082 |
| | Mean B | 0.2082 | 0.2807 | 0.2807 |
| | Δ (A–B) | +0.1488 | +0.0763 | -0.0725 |
| | t-stat | 11.61 | 7.67 | -3.74 |
| | p-value | 0.0003 | 0.0016 | 0.0202 |
| | Significance | *** | ** | * |

***Table S7**. **Full GNN performance under graph-structure and feature-ablation conditions.*** *Test-set performance of the GNN across STRING, degree-preserving shuffled, and fully random graph conditions, evaluated with and without network-derived node features. Results are reported as mean ± standard deviation across five gene-level cross-validation folds.MSE denotes the weighted mean squared error used as the training loss.*

| **Breast - with network features** | | | |
|---|---|---|---|
| **Metric** | **STRING Test** | **Degree-preserving Test** | **Random Test** |
| **MSE** | 0.2003 ± 0.0112 | 0.2248 ± 0.0053 | 0.2313 ± 0.0131 |
| **Pearson** | 0.5047 ± 0.0528 | 0.4170 ± 0.0336 | 0.4227 ± 0.0335 |

| | | | |
|---|---|---|---|
| **Spearman** | 0.4867 ± 0.0510 | 0.3859 ± 0.0342 | 0.4003 ± 0.0261 |
| **Balanced Accuracy** | 0.6734 ± 0.0231 | 0.6260 ± 0.0192 | 0.6321 ± 0.0176 |
| **F1 macro** | 0.6727 ± 0.0241 | 0.6253 ± 0.0204 | 0.6317 ± 0.0181 |
| **F1 weighted** | 0.6764 ± 0.0246 | 0.6298 ± 0.0200 | 0.6361 ± 0.0165 |
| **MCC** | 0.3524 ± 0.0430 | 0.2562 ± 0.0367 | 0.2680 ± 0.0321 |
| **Breast - without network features** | | | |
| **MSE** | 0.1999 ± 0.0136 | 0.2373 ± 0.0067 | 0.2783 ± 0.0081 |
| **Pearson** | 0.5163 ± 0.0451 | 0.3779 ± 0.0359 | 0.2331 ± 0.0423 |
| **Spearman** | 0.5125 ± 0.0402 | 0.3742 ± 0.0371 | 0.2412 ± 0.0374 |
| **Balanced Accuracy** | 0.6809 ± 0.0265 | 0.6209 ± 0.0154 | 0.5841 ± 0.0085 |
| **F1 macro** | 0.6788 ± 0.0298 | 0.6192 ± 0.0160 | 0.5812 ± 0.0101 |
| **F1 weighted** | 0.6827 ± 0.0297 | 0.6245 ± 0.0172 | 0.5868 ± 0.0131 |
| **MCC** | 0.3710 ± 0.0450 | 0.2490 ± 0.0312 | 0.1738 ± 0.0193 |
| **Lung - with network features** | | | |
| **MSE** | 0.1774 ± 0.0070 | 0.2013 ± 0.0121 | 0.2058 ± 0.0208 |
| **Pearson** | 0.5343 ± 0.0243 | 0.4473 ± 0.0338 | 0.4399 ± 0.0507 |
| **Spearman** | 0.5105 ± 0.0321 | 0.4122 ± 0.0331 | 0.4138 ± 0.0492 |
| **Balanced Accuracy** | 0.6879 ± 0.0155 | 0.6430 ± 0.0136 | 0.6461 ± 0.0159 |
| **F1 macro** | 0.6871 ± 0.0151 | 0.6422 ± 0.0144 | 0.6458 ± 0.0161 |
| **F1 weighted** | 0.6891 ± 0.0152 | 0.6449 ± 0.0146 | 0.6486 ± 0.0170 |
| **MCC** | 0.3756 ± 0.0307 | 0.2871 ± 0.0275 | 0.2932 ± 0.0333 |
| **Lung - without network features** | | | |
| **MSE** | 0.1845 ± 0.0112 | 0.2205 ± 0.0108 | 0.2774 ± 0.0660 |
| **Pearson** | 0.5321 ± 0.0155 | 0.3866 ± 0.0196 | 0.2498 ± 0.0517 |
| **Spearman** | 0.5237 ± 0.0214 | 0.3771 ± 0.0182 | 0.2483 ± 0.0358 |
| **Balanced Accuracy** | 0.6946 ± 0.0126 | 0.6262 ± 0.0111 | 0.5811 ± 0.0134 |
| **F1 macro** | 0.6936 ± 0.0122 | 0.6256 ± 0.0120 | 0.5775 ± 0.0160 |
| **F1 weighted** | 0.6960 ± 0.0123 | 0.6303 ± 0.0098 | 0.5828 ± 0.0188 |
| **MCC** | 0.3913 ± 0.0213 | 0.2568 ± 0.0193 | 0.1674 ± 0.0277 |
| **Large Intestine - with network features** | | | |
| **MSE** | 0.1867 ± 0.0082 | 0.2139 ± 0.0074 | 0.2274 ± 0.0154 |
| **Pearson** | 0.5101 ± 0.0099 | 0.4182 ± 0.0308 | 0.3805 ± 0.0387 |
| **Spearman** | 0.4922 ± 0.0100 | 0.3881 ± 0.0342 | 0.3493 ± 0.0408 |

| | | | |
|---|---|---|---|
| **Balanced Accuracy** | 0.6777 ± 0.0045 | 0.6299 ± 0.0124 | 0.6035 ± 0.0155 |
| **F1 macro** | 0.6778 ± 0.0047 | 0.6298 ± 0.0129 | 0.5973 ± 0.0205 |
| **F1 weighted** | 0.6808 ± 0.0055 | 0.6339 ± 0.0138 | 0.6046 ± 0.0199 |
| **MCC** | 0.3570 ± 0.0092 | 0.2627 ± 0.0253 | 0.2198 ± 0.0288 |
| **Large Intestine - without network features** | | | |
| **MSE** | 0.1890 ± 0.0124 | 0.2228 ± 0.0228 | 0.2584 ± 0.0153 |
| **Pearson** | 0.5172 ± 0.0364 | 0.4040 ± 0.0436 | 0.2578 ± 0.0448 |
| **Spearman** | 0.5129 ± 0.0248 | 0.3899 ± 0.0378 | 0.2491 ± 0.0406 |
| **Balanced Accuracy** | 0.6781 ± 0.0179 | 0.6244 ± 0.0128 | 0.5791 ± 0.0156 |
| **F1 macro** | 0.6769 ± 0.0197 | 0.6240 ± 0.0128 | 0.5749 ± 0.0155 |
| **F1 weighted** | 0.6822 ± 0.0174 | 0.6290 ± 0.0135 | 0.5827 ± 0.0159 |
| **MCC** | 0.3701 ± 0.0245 | 0.2539 ± 0.0278 | 0.1665 ± 0.0337 |

***Table S8**. **Impact of graph topology on GNN performance.** Comparison of GNN performance across three graph structures: biologically derived STRING network, degree-preserving shuffled network, and fully random network. Performance is evaluated on test-set weighted MSE (Loss) and MCC across 5 cross-validation folds, with statistical significance assessed using paired t-tests.*

| **Breast** | | | | |
|---|---|---|---|---|
| **Metric** | **Statistic** | **STRING vs Shuffled** | **STRING vs Random** | **Shuffled vs Random** |
| **MSE** | **Mean (A)** | 0.2003 | 0.2003 | 0.2248 |
| | **Mean (B)** | 0.2248 | 0.2313 | 0.2313 |
| | **Δ** | -0.0245 | -0.0310 | -0.0065 |
| | **t-stat** | -4.12 | -3.25 | -1.10 |
| | **p-value** | 0.014 | 0.031 | 0.330 |
| | **Significance** | * | * | n.s. |
| **MCC** | **Mean (A)** | 0.3524 | 0.3524 | 0.2562 |
| | **Mean (B)** | 0.2562 | 0.2680 | 0.2680 |
| | **Δ** | +0.0962 | +0.0844 | -0.0118 |
| | **t-stat** | 3.01 | 2.80 | -0.42 |
| | **p-value** | 0.038 | 0.049 | 0.690 |

| | | | | |
|---|---|---|---|---|
| | **Significance** | * | * | n.s. |
| | | **Lung** | | |
| **MSE** | **Mean (A)** | 0.1774 | 0.1774 | 0.2013 |
| | **Mean (B)** | 0.2013 | 0.2058 | 0.2058 |
| | **Δ** | -0.0239 | -0.0284 | -0.0045 |
| | **t-stat** | -3.80 | -2.90 | -0.95 |
| | **p-value** | 0.019 | 0.044 | 0.390 |
| | **Significance** | * | * | n.s. |
| **MCC** | **Mean (A)** | 0.3756 | 0.3756 | 0.2871 |
| | **Mean (B)** | 0.2871 | 0.2932 | 0.2932 |
| | **Δ** | +0.0885 | +0.0824 | -0.0061 |
| | **t-stat** | 3.45 | 3.10 | -0.30 |
| | **p-value** | 0.025 | 0.035 | 0.780 |
| | **Significance** | * | * | n.s. |
| | | **Large Intestine** | | |
| **MSE** | **Mean (STRING)** | 0.1867 | 0.1867 | 0.2139 |
| | **Mean (Other)** | 0.2139 | 0.2274 | 0.2274 |
| | **Δ** | -0.0272 | -0.0407 | -0.0135 |
| | **t-stat** | -4.50 | -5.10 | -1.40 |
| | **p-value** | 0.010 | 0.007 | 0.240 |
| | **Significance** | * | ** | n.s. |
| **MCC** | **Mean (STRING)** | 0.3570 | 0.3570 | 0.2627 |
| | **Mean (Other)** | 0.2627 | 0.2198 | 0.2198 |
| | **Δ** | +0.0943 | +0.1372 | -0.0429 |
| | **t-stat** | 4.00 | 4.60 | -1.20 |

| | | | | |
|---|---|---|---|---|
| | **p-value** | 0.016 | 0.009 | 0.310 |
| | **Significance** | * | ** | n.s. |

***Table S9*. *Effect of network structure on GNN performance across tissues.*** *Comparison of Graph Neural Network (GNN) models using full network connectivity versus models without network features ("no-net") across three graph conditions: biologically derived STRING network, degree-preserving shuffled network, and fully random network. Performance is evaluated using weighted mean squared error (Loss) and Matthews correlation coefficient (MCC) on the test set across 5 cross-validation folds. Statistical significance was assessed using a two-sided paired t-test on per-fold results.*

| **Breast** (with vs without-network features) | | | | |
|---|---|---|---|---|
| **Metric** | **Statistic** | **STRING** | **Degree-preserving** | **Random** |
| **MSE** | Mean (A) | 0.2003 | 0.2248 | 0.2313 |
| | Mean (B) | 0.1999 | 0.2373 | 0.2783 |
| | Δ | +0.0005 | -0.0125 | -0.0470 |
| | t-stat | 0.06 | -3.06 | -4.98 |
| | p-value | 0.956 | 0.038 | 0.0076 |
| | Significance | n.s. | * | ** |
| **MCC** | Mean (A) | 0.3524 | 0.2562 | 0.2680 |
| | Mean (B) | 0.3710 | 0.2490 | 0.1738 |
| | Δ | -0.0186 | +0.0072 | +0.0942 |
| | t-stat | -0.60 | 0.33 | 4.73 |
| | p-value | 0.579 | 0.761 | 0.0091 |
| | Significance | n.s. | n.s. | ** |
| **Lung** (with vs without-network features) | | | | |
| **MSE** | Mean (A) | 0.1774 | 0.2013 | 0.2058 |
| | Mean (B) | 0.1845 | 0.2205 | 0.2774 |
| | Δ | -0.0071 | -0.0192 | -0.0716 |

| | | | | |
|---|---|---|---|---|
| | t-stat | -1.12 | -3.03 | -2.17 |
| | p-value | 0.325 | 0.0388 | 0.0957 |
| | Significance | n.s. | * | n.s. |
| **MCC** | Mean (A) | 0.3756 | 0.2871 | 0.2932 |
| | Mean (B) | 0.3913 | 0.2568 | 0.1674 |
| | Δ | -0.0157 | +0.0303 | +0.1258 |
| | t-stat | -0.91 | 3.07 | 5.12 |
| | p-value | 0.415 | 0.0372 | 0.0069 |
| | Significance | n.s. | * | ** |
| **Large Intestine**<br>(with vs without-network features) | | | | |
| **MSE** | Mean (A) | 0.1867 | 0.2139 | 0.2274 |
| | Mean (B) | 0.1890 | 0.2228 | 0.2584 |
| | Δ | -0.0023 | -0.0089 | -0.0310 |
| | t-stat | -0.54 | -0.66 | -8.39 |
| | p-value | 0.619 | 0.543 | 0.0011 |
| | Significance | n.s. | n.s. | ** |
| **MCC** | Mean (A) | 0.3570 | 0.2627 | 0.2198 |
| | Mean (B) | 0.3701 | 0.2539 | 0.1665 |
| | Δ | -0.0131 | +0.0088 | +0.0533 |
| | t-stat | -1.00 | 0.39 | 5.87 |
| | p-value | 0.373 | 0.714 | 0.0042 |
| | Significance | n.s. | n.s. | ** |

***Table S10*. Comparison of GNN and Graph Transformer (GPS) architectures.** *Performance comparison between Graph Neural Networks (GNN) and Graph Transformer models (GPS) across tissues, evaluated using weighted MSE (Loss) and Matthews correlation coefficient (MCC) on the test set across 5 cross-validation folds. Statistical significance was assessed using a two-sided paired t-test.*

| **Metric** | **Statistic** | **GNN vs GPS (Breast)** | **GNN vs GPS (Lung)** | **GNN vs GPS (Large intestine)** |
|---|---|---|---|---|
| Loss | Mean (GNN) | 0.2003 | 0.1774 | 0.1867 |
| | Mean (GPS) | 0.2331 | 0.1936 | 0.2072 |
| | Δ (GNN − GPS) | -0.0328 | -0.0163 | -0.0205 |
| | t-stat | -8.02 | -2.52 | -6.87 |
| | p-value | 0.0013 | 0.0653 | 0.0024 |
| | Sig | ** | n.s. | ** |
| MCC | Mean (GNN) | 0.3524 | 0.3756 | 0.3570 |
| | Mean (GPS) | 0.2757 | 0.3394 | 0.3068 |
| | Δ (GNN − GPS) | +0.0767 | +0.0362 | +0.0502 |
| | t-stat | 4.93 | 3.99 | 3.08 |
| | p-value | 0.0079 | 0.0162 | 0.0370 |
| | Sig | ** | * | * |
